# Transcranial Photobiomodulation with Near-Infrared Light from Childhood to Elderliness: Simulation of Dosimetry

**DOI:** 10.1101/2020.02.06.937821

**Authors:** Yaoshen Yuan, Paolo Cassano, Matthew Pias, Qianqian Fang

## Abstract

**Significance:** Major depressive disorder (MDD) affects over 40 million US adults in their lifetimes. Transcranial photobiomodulation (t-PBM) has been shown to be effective in treating MDD, but the current treatment dosage does not account for anatomical head and brain changes due to aging.

**Aim:** We study effective t-PBM dosage and its variations across age groups using state-of-the-art Monte Caxrlo (MC) simulations and age-dependent brain atlases ranging between 5 to 85 years of age.

**Approach:** Age-dependent brain models are derived from 18 MRI brain atlases. Two extracranial source positions, F3-F4 and Fp1-Fpz-Fp2 in the EEG 10-20 system, are simulated at five selected wavelengths and energy depositions at two MDD-relevant cortical regions – dorsolateral prefrontal cortex (dlPFC) and ventromedial prefrontal cortex (vmPFC) – are quantified.

**Results:** An overall decrease of energy deposition was found with increasing age. A strong negative correlation between the thickness of extra-cerebral tissues (ECT) and energy deposition, suggesting that increasing ECT thickness over age is primarily responsible for reduced energy delivery. The F3-F4 position appears to be more efficient in reaching dlPFC compared to treating vmPFC via the Fp1-Fpz-Fp2 position.

**Conclusion:** Quantitative simulations revealed age-dependent light delivery across the lifespan of human brains, suggesting the needs for personalized and age-adaptive t-PBM treatment planning.

## 1 Introduction

According to National Institute of Health,^1^ the estimated lifetime prevalence of major depressive disorder (MDD) in the United States comprises of over 13% of the population. MDD can develop at any age, and is considered the leading cause of disability in the US for individuals between the ages of 15 and 44.^2–4^ The two most commonly used treatments for MDD are antidepressants (87.0%) and psychotherapy (23.2%).^5^ Several known challenges have been faced by these treatment approaches: 1) frequent relapses of the cognitive therapy^6^ and 2) burdensome side-effects of antidepressant medication.^7^ Furthermore, many patients prefer to self-manage, which leads to the low treatment rates.^8^ Therefore, new, effective, safe, and easy-to-administer treatment methods are needed to battle MDD.

Photobiomodulation (PBM) is a near-infrared (NIR) light-based therapy technique and has shown therapeutic effectiveness for various neuropsychiatric disorders, including MDD.^9–11^ The transcranial PBM (t-PBM) technique delivers NIR light through the scalp and skull.^12–14^ Due to the penetration depth of NIR light in human tissues, clinically effective light dosages can be delivered to the disease-responsible brain regions without damaging superficial tissues. While the molecular mechanisms of PBM remain a topic of active research, some studies report that treatment effects may derive from the excitation of a mitochondrial chromophore – cytochrome c oxidase (CCO) – at the NIR spectra,^14^ stimulating the mitochondrial respiratory chain and increasing ATP production.^10, 11^ The concurrent upshot of reactive oxygen species (ROS) may trigger cytoprotective and anti-oxidation pathways within the cell, with effects potentially lasting on the scale of days to weeks.^15^ A wide range of studies, in both animal models and humans, have shown that PBM causes minimal or no adverse effects while producing therapeutic effects.^10, 16, 17^

Although MDD has a broad age of onset,^2, 4^ most previously published studies have focused on t-PBM treatments in only middle-aged adult brain models.^18^ However, personalization of treatments is key to increasing success rate and tolerability; therefore our interest is in developing precise PBM treatment strategies adapted for individual patients. One of the main factors impacting t-PBM light dosage is the thickness of extra-cerebral tissues (ECT), including both skull and scalp.^19–21^ Therefore, a quantitative analysis on how brain development and senescence could impact the effective dosage in a t-PBM treatment can provide valuable guidelines for clinicians to optimize their procedures and maximize treatment efficacy and tolerability.

To capture the variations of anatomical features among age groups, we have to first create anatomically appropriate brain/full-head models, including skin/skull/brain 3-D shapes and thicknesses. Fortunately, a number of recent studies have published comprehensive MRI atlases outlining the development of human brains from infants to elders.^22–25^ In addition, several groups, including our own, have developed sophisticated brain segmentation and meshing pipelines to convert neuroanatomical scans into high quality multi-layered brain models. These resources make it possible to quantitatively investigate how the development and senescence of the human brain influences light penetration at different stages of life.

In addition, advanced photon transport models must be used to accurately account for the complex light-tissue interactions during t-PBM procedures. In this study, we applied the Monte Carlo (MC) method – a stochastic solver for the radiative transfer equation (RTE) – which is widely considered the gold standard for light modeling in complex tissues.^26^ While alternative models, such as the diffusion equation (DE), are dramatically faster and applicable to many types of human tissues,^27, 28^ for brain tissues, DE is known to produce erroneous solutions due to the presence of low-scattering media such as air cavities and cerebrospinal fluid (CSF).^29, 30^ The MC method solves the RTE rigorously by simulating large numbers of photons following a set of known probability models derived from physics.^31^ The only major limitation is that MC methods are computationally expensive. To improve computational efficiency, we applied our widely disseminated hardware-accelerated Monte Carlo modeling platform – Monte Carlo eXtreme (MCX).^32^ This tool can shorten the simulation runtime by several hundreds fold compared to conventional CPU based simulations.^32, 33^

The remainder of the paper is organized as follows. In the Materials and Methods section, we detail the preprocessing steps to create 4-layer head segmentations from the neurodevelopmental MRI brain atlas library.^22^ We also report the steps to obtain brain parcellations and the placement of light source positions. In the Results section, the simulated energy depositions for 2 t-PBM source placements, 5 selected wavelengths on 18 selected brain/head atlases, ranging between 5 and 89 years of age are reported. In the Discussion section, we highlight the findings regarding the efficiency of different wavelengths, the energy deposition and the exposure duration in the wide span of age groups. In addition, we also correlate our findings with the anatomical changes associated with the brain development and senescence.

## 2 Materials and methods

### 2.1 Creating multi-layer head models from MRI brain atlas library

Brain segmentations are created by processing the Neurodevelopmental MRI database.^22–24^ In this study, we select a total of 18 age groups, ranging from 5 through 89 years of age. Specifically, the average atlas for 5, 10, 14, 18, 20-24, 25-29, 30-34, 35-39, 40-44, 45-49, 50-54, 55-59, 60-64, 65-69, 70-74, 75-79, 80-84, and 85-89 age groups are used in our study. For each atlas, a 4-layer full-head segmentation is created, including the white matter (WM), gray matter (GM), CSF and ECT. An additional air cavity segmentation is created to properly model light propagation inside the nasal and pharyngeal cavities. As a result, a total of 5 tissue labels are considered. Three sample segmented brain volumes at 5, 40-44, 85-89 years of age are shown in Figs. 1(a)-1(c).

**Fig 1.**
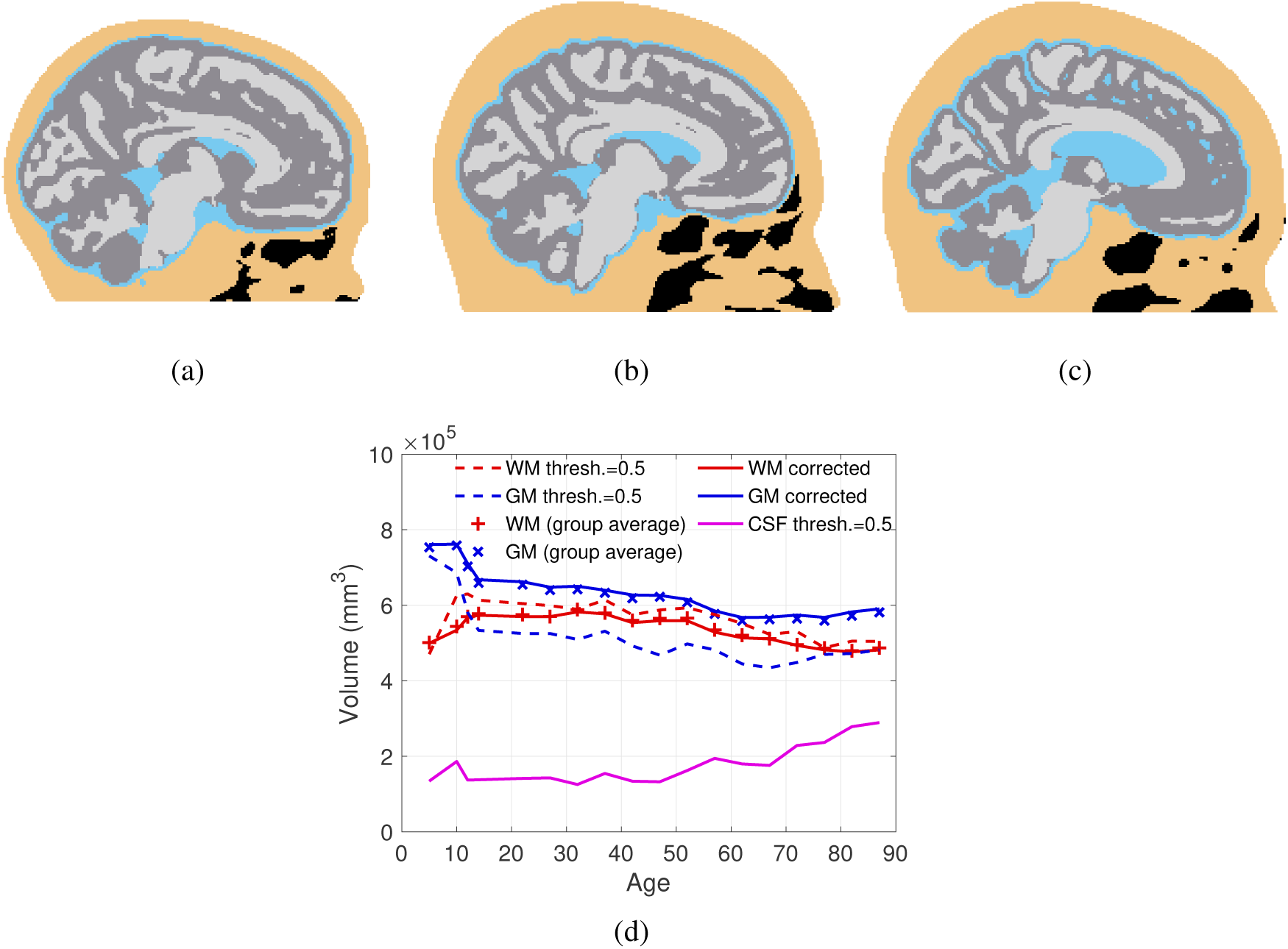
Sample segmented brain models for 5, 40-44, 85-89 years old are shown in (a), (b) and (c) respectively to visualize brain development. Black regions represent air cavities. In (d) we show the white-matter (WM), gray-matter (GM) and cerebrospinal fluid (CSF) volumes before (dashed-lines) and after (solid-lines) adaptive thresholding. The group-average WM and GM volumes derived from the source population are based on Refs.^23, 24^

The brain atlas probabilistic tissue segmentations (PTS) provided in the Neurodevelopmental MRI database were derived from averaging subjects in each age group. The GM/WM volumes directly calculated from the atlas PTS volumes using a simple threshold show discrepancies compared to the GM/WM volumes estimated from the original group-based data published by the same authors.^23, 24^ We believe that this discrepancy results from the averaging and non-linear effects of the atlas creation process.

To correct for this discrepancy, an adaptive threshold (*T* ∈ [0,1]) is applied to the PTSs of WM, GM and CSF of all atlases. As shown in Fig. 1(d), before this correction, a uniform threshold *T* = 0.5 of the atlas segmentation (dashed lines) appears to underestimate the GM volume and overestimate the WM volume along age compared to the previously reported averaged volumes of the population^23, 24^ from which the atlases are derived. To reduce this artifact, we dynamically estimate a threshold for GM/WM to match the tissue volumes to the population-derived estimations. The corrected GM/WM volumes (solid lines) over age are shown in Fig. 1(d). in comparison, such discrepancies for the CSF layer are relatively small compared to previous studies.^23, 24, 34, 35^ For simplicity, a uniform threshold *T*_C*SF*_ = 0.5 is applied to T1-weighted (T1w) CSF PTS.

To obtain the exterior surface of the ECT layer, i.e. the scalp surface, we use FSL and the `betsurf` add-on to process the T1w MRI images provided by the atlas database.^23, 36^ Note that the FSL pipeline for segmenting ECT tissues has been validated in adult brains/heads. For young age children, only limited studies have been reported.^37–39^ Therefore, our results for young children are intended for qualitative assessment only.

### 2.2 Segmentations of air cavities in the brain atlases

It is important to note that the presence of air cavities in the brain atlases has significant impact to light dosimetry due to the low absorption of air. However, most neuroanatomical analysis tools do not have the ability to automatically extract these cavities. Here, we use a combination of manual segmentation and clustering analysis to extract various head cavities.

The frontal sinus and sphenoid sinus are manually segmented using the T1w MRI images and ITK-SNAP.^40^ To segment the nasal and pharyngeal cavities, we apply the *k*-means algorithm to the raw T1w MRI images.^41^ A total of three clusters are segmented for the ECT layer below Fpz point and the cluster with the lowest intensity is used as the air-cavities. The eyes and the spine also appear as low intensity regions in T1w images. A manual flood-filling operation is performed to identify these regions. Examples of the created multi-layer brain anatomies are shown in Fig. 2(a).

**Fig 2.**
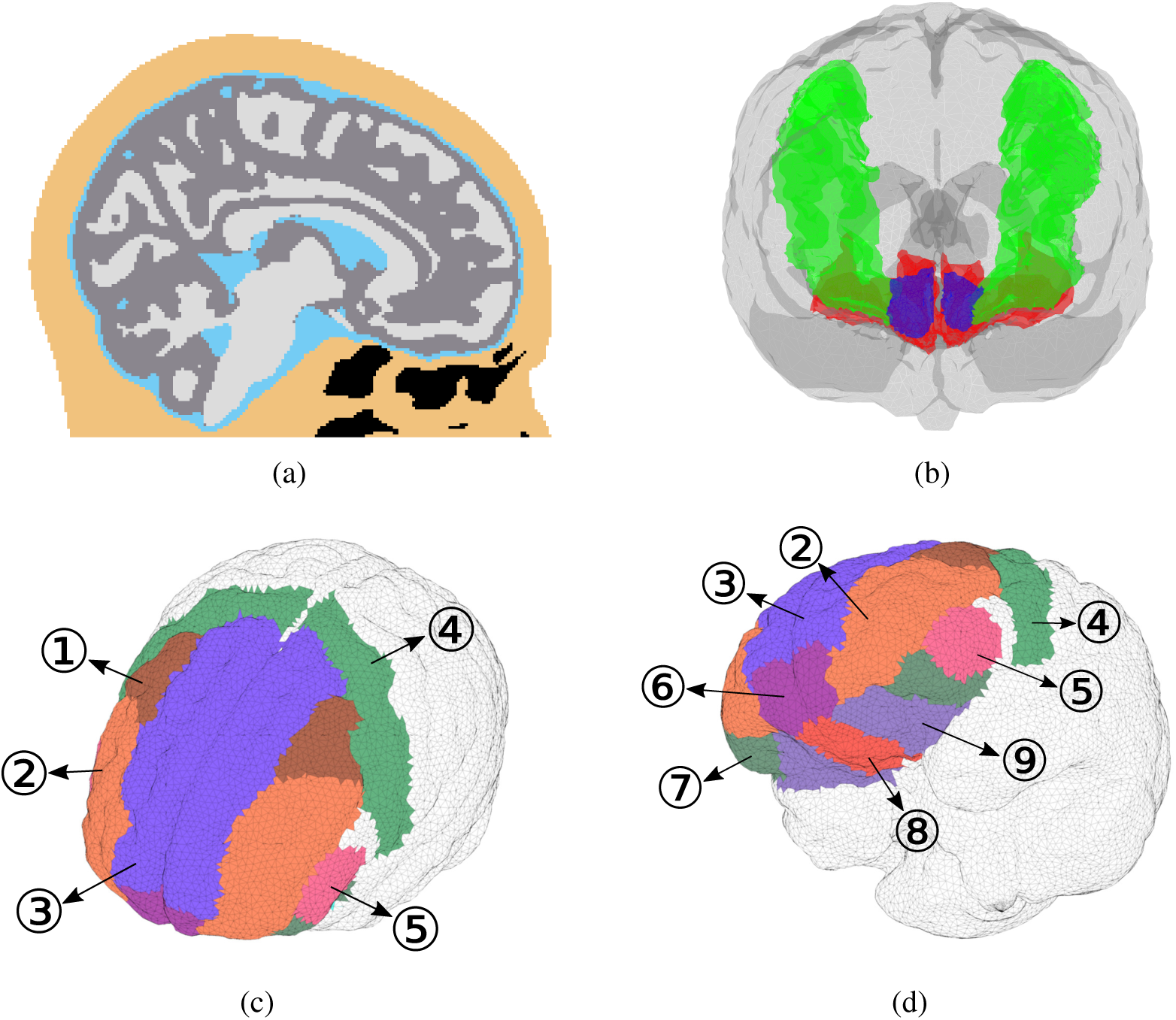
Sample brain segmentation and regions-of-interest. (a) 18-0 years old 4-layer model, including WM, GM, CSF, extra-cerebral tissues (ECT). Black regions indicate air cavities. In (b), green regions are dlPFC and red regions are vmPFC. The frontal pole is indicated by blue color. In (c)-(d), we labeled all parcellations (bilateral) used in this study, including 1-caudal middle frontal gyrus, 2-rostral middle frontal gyrus, 3-superior frontal gyrus, 4-precentral gyrus, 5-pars triangularis, 6-frontal pole, 7-pars orbitalis, 8-medial orbitofrontal cortex, 9-lateral orbitofrontal cortex.

### 2.3 Brain parcellations and target regions

Similar to our previous study,^18^ our primary interest here is to develop effective PBM treatment approaches for emotion regulation and depression. Thus, our main focuses in this dosimetry study are the dlPFC and vmPFC regions (Fig. 2(b)), both involved in the emotion regulation circuitry. However, our approach is general and can be used to characterize all brain functional regions. We used the MarsAtlas parcellation in our previous study.^18, 42^ This parcellation of the vmPFC region includes the frontal pole (Brodmann Area 10) while several other studies did not.^43–46^ This may potentially cause a discrepancy in calculations of energy deposition in vmPFC. In this study, we consider both definitions by selecting and merging sub-regions from the Desikan-Killiany-Tourville (DKT) parcellation^47^ that best comprises the vmPFC regions in either definition. The brain DKT parcellation is obtained by using the `recon-all` workflow provided in FreeSurfer v6.0,^48, 49^ see Figs. 2(c)-2(d). The FreeSurfer brain parcellation workflow has been validated in previous publications for ages above 3-years of age.^45, 50^

In this study, dlPFC is considered to cover two regions in the DKT parcellation – caudal middle frontal gyrus and rostral middle frontal gyrus (Labels 1 and 2 in Fig. 2(c)) according to Ref.^51^ The superior frontal gyrus is excluded because it is not relevant to t-PBM. As mentioned earlier, a general consensus on the boundary of the vmPFC has not been reached. In previous literature, vmPFC is either defined as the combination of 1) medial orbitofrontal cortex and lateral orbitofrontal cortex,^45, 46^ i.e. Labels 8 and 9 in Fig. 2(d) or 2) frontal pole, medial orbitofrontal cortex and lateral orbitofrontal cortex, i.e. Labels 6, 8 and 9 in Fig. 2(d), referred to as the extended vmPFC^52, 53^ hereinafter. Furthermore, due to the slight mismatch between the FreeSurfer parcellations and the GM in our 4-layer segmentation, the final parcellations are defined as the intersection between the two models.

### 2.4 Light source positions

In this study we focus on characterizing the transcranial PBM (t-PBM) treatment strategy. A pair of transcranial source configurations^18^ are investigated: 1) **F3-F4**: two light-emitting-diode (LED) array sources are placed above and centered, respectively, at the F3 and F4 (10-20 EEG positions); 2) **Fp1-Fpz-Fp2:** one LED-array is placed on the forehead and between the Fp1-Fpz-Fp2 positions (referred to as the Fpz position hereinafter). The definitions of these source positions and orientations are similar to,^18^ with the exception that the F3-F4 source arrays are rotated around 90 degrees, as shown in Fig. 3, to better align with the dlPFC region and maximize light delivery. The simulated LED-arrays follow the dimensions and parameters of an available PBM source (Omnilux New-U, Photomedex, Horsham, PA).

**Fig 3.**
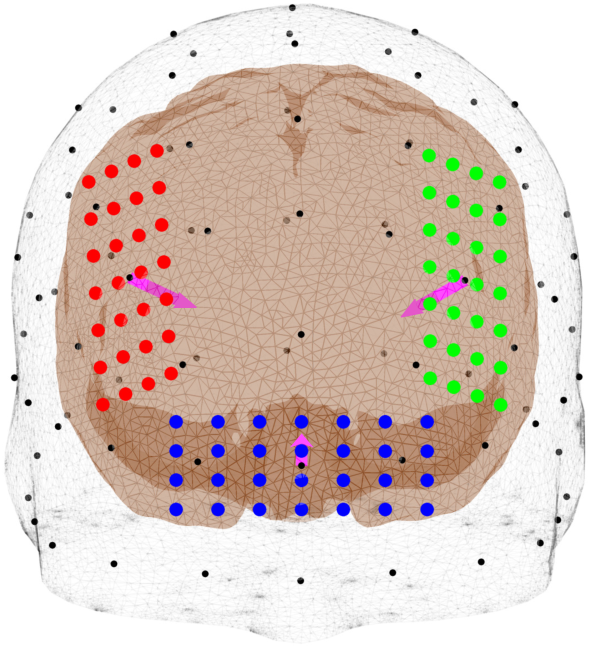
Illustrations of source positions. Three extracranial source positions are shown – F3 (green), F4 (red), and Fpz (blue). Each source is an array of light-emitting diodes represented by colored dots. The magenta arrow represents the source direction.

### 2.5 Simulation settings

In our study, a total of 15 simulations, a combination of 3 source positions and 5 wavelengths (670 nm, 810 nm, 850 nm, 980 nm and 1064 nm), are performed for each segmented brain atlas using our GPU-accelerated photon transport simulator (MCX).^32, 33^ The optical properties of each layer (WM, GM and CSF) are identical to those from our earlier study,^18^ with the exception that the optical properties of the ECT layer (Table 1) are derived using the weighted average of the properties for fat, muscle, skin tissue and skull; the weights are derived from volume fractions of the respective tissue types in the Colin27 atlas.^54, 55^ For simplicity, we assume the optical properties for each tissue type are independent of age. In all simulations, a total of 10^9^ photons are launched. To reduce the effect of shot-noise in regions distal to the light source, an adaptive non-local mean filter is applied.^56^ All atlases share the same resolution of 1 × 1 × 1 mm^3^ isotropic voxels. Normalized average energy deposition (J/cm^3^), as described in,^18^ is then computed for each atlas and compared across different age groups. The only exception is that peak fluence (99^th^ percentile of the target) is adopted for computing the exposure duration for one treatment session.

**Table 1.**
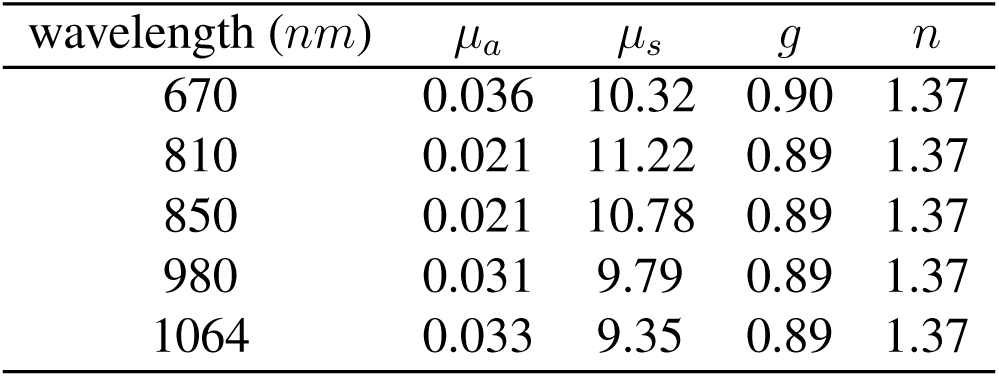
The estimated optical properties of the extra-cerebral tissues (ECT) layer at 5 wavelengths. *µ*_*a*_ – absorption coefficient (mm^*−*1^), *µ*_*s*_ – scattering coefficient (mm^*−*1^), *g* – anisotropy, and *n* – refractive index.

| wavelength (nm) | $\mu_a$ | $\mu_s$ | $g$ | $n$ |
| --- | --- | --- | --- | --- |
| 670 | 0.036 | 10.32 | 0.90 | 1.37 |
| 810 | 0.021 | 11.22 | 0.89 | 1.37 |
| 850 | 0.021 | 10.78 | 0.89 | 1.37 |
| 980 | 0.031 | 9.79 | 0.89 | 1.37 |
| 1064 | 0.033 | 9.35 | 0.89 | 1.37 |

### 2.6 Extra-cerebral tissues (ECT) thickness estimation

To further understand the age-dependency of t-PBM light dosage, we also compute the thickness of the ECT layer. In Fig. 4, three ECT regions for F3 (green), F4 (red) and Fpz (blue) positions are created for computing the ECT thickness. The average ECT thicknesses are estimated by 1) using the “Iso2Mesh” toolbox^57^ to create meshes for the inner and outer surfaces of the ECT layer for each atlas, 2) projecting the illuminated areas from the outer surface inward along the normal direction to generate a truncated volume, and 3) computing the average thickness by dividing the enclosed volume by the area of the illuminated area.

**Fig 4.**
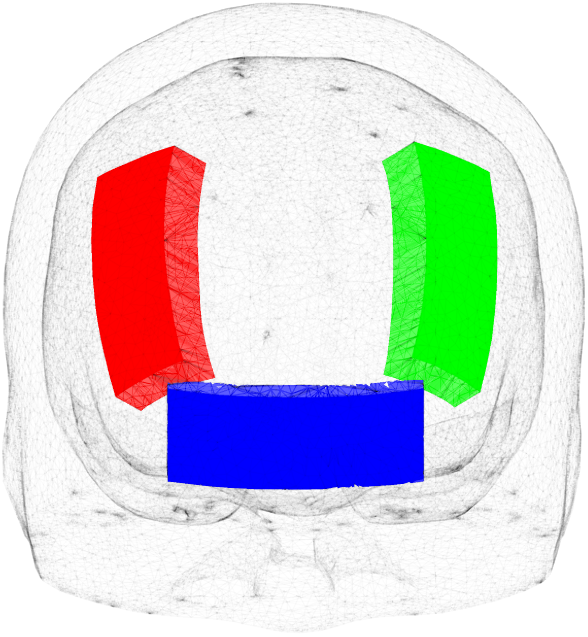
The extra-cerebral tissues (ECT) regions used to compute the ECT thickness under F3 (green), F4 (red) and Fpz (blue) positions.

## 3 Results

### 3.1 Photon dosimetry assessment

Sample sagittal energy deposition maps are shown in Figs. 5(a)-5(c) to demonstrate qualitatively the changes of light distributions over age. The selected plots are generated using an Fpz source at 810 nm for 3 different age groups – 5, 40-44, 85-89. In Fig. 6, we summarize the age-dependent average energy depositions in dlPFC and vmPFC using F3-F4 and Fpz positions respectively for 5 selected wavelengths. Similar to our previous findings,^18^ 810 nm illumination appears to provide the highest energy deposition across a wide range of age spans, followed by 1064 nm and 850 nm. In addition, a strong linear correlation on this log-scaled plot is found between wavelengths. For extended vmPFC (not included in Fig. 6), the same conclusions can be drawn. For simplicity, hereinafter, we only report results at 810 nm. In Fig. 7, we show the top-5 brain parcellations that have received the highest average energy deposition. The parcellations in the legend are roughly ranked in descending order based on energy deposition, although such orders may vary slightly across age groups. The numbers in parentheses correspond to the numbering shown in Figs. 2(c)-2(d).

**Fig 5.**
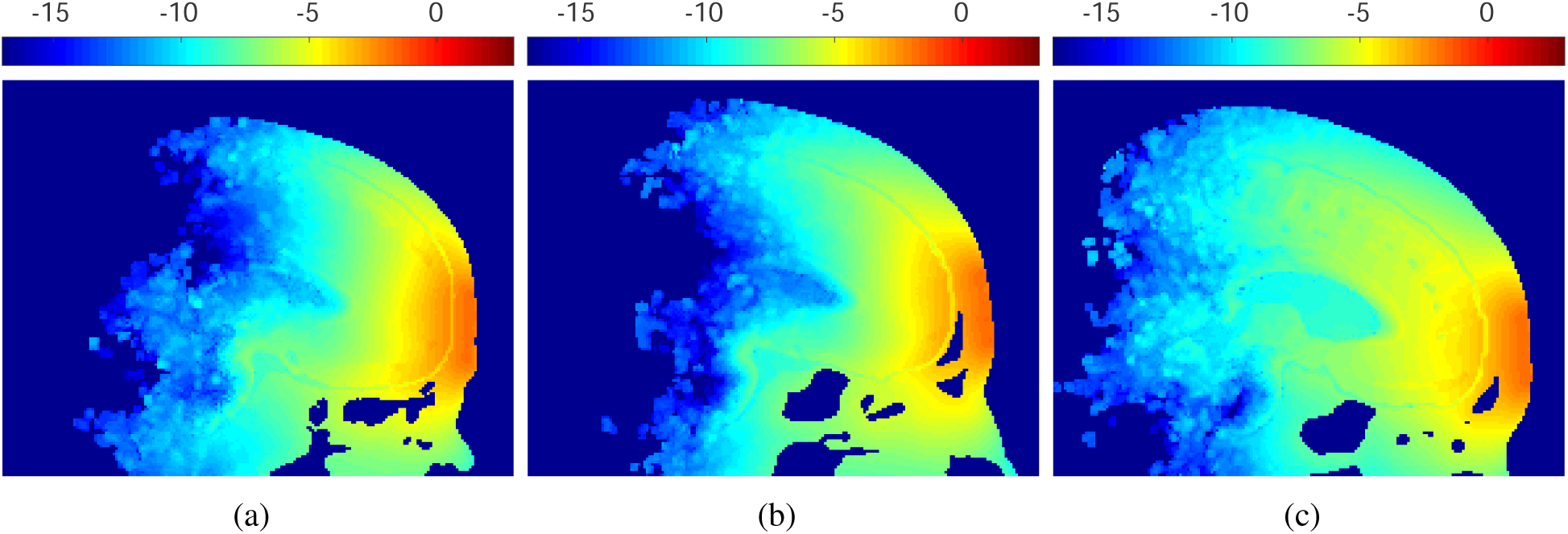
Sample energy distributions. From (a) to (c), we show energy deposition (J/cm^3^) in log_10_ scale for 5, 40-44, 85-89 years old, respectively. Only Fpz source position and 810 nm results are shown here.

**Fig 6.**
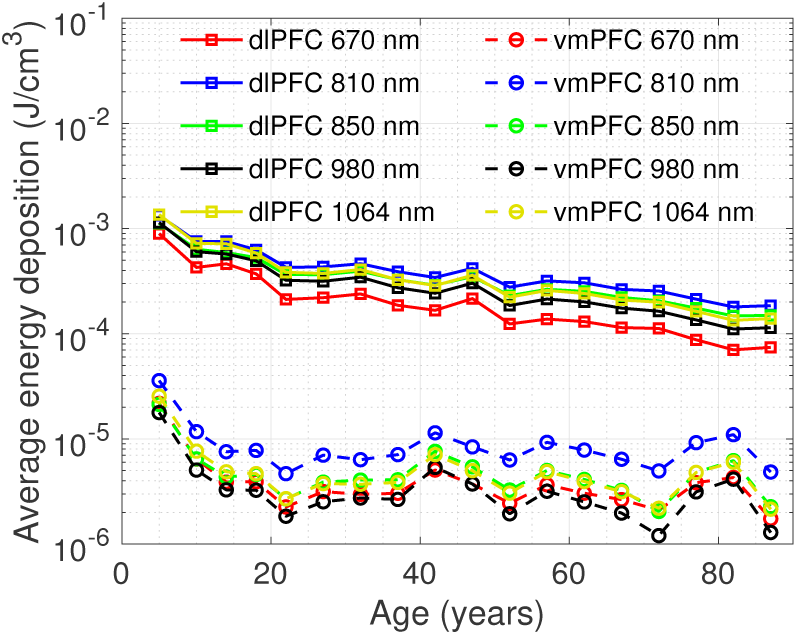
Average energy deposition for 5 wavelengths across age. The plots for various wavelengths are shown in different colors. The solid line with square markers and the dashed line with circular markers indicate dlPFC using F3-F4 position and vmPFC using Fpz position respectively.

**Fig 7.**
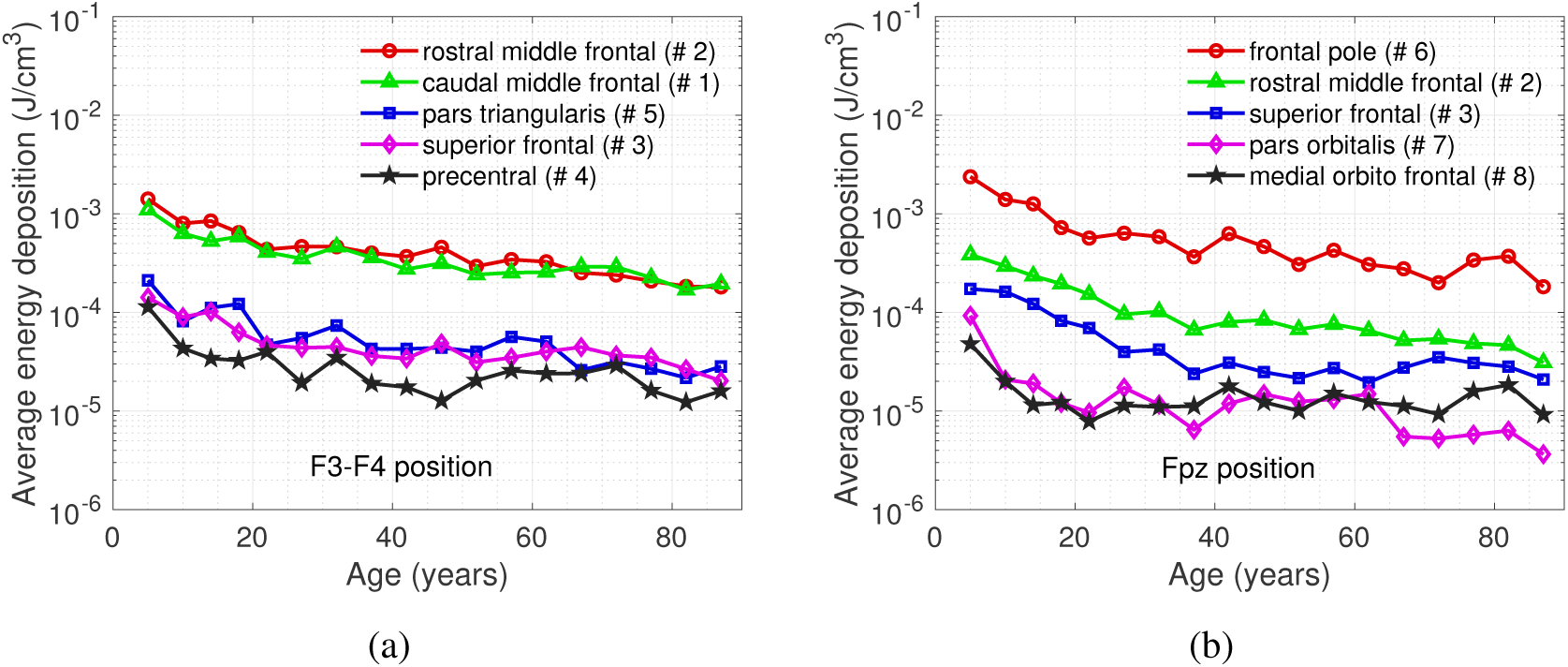
Average energy deposition variations across age. We show results for (a) F3-F4 and (b) Fpz source positions respectively. Each plot includes the parcellations with top-5 average energy deposition in descending order from top to bottom in the legend. The numbers in the parenthesis correspond to the parcellation labels in Figs. 2(c) and 2(d). Only results from 810 nm are shown.

In Fig. 8(a), we show the average energy deposition for dlPFC and vmPFC using both source positions. In Fig. 8(b), we show the exposure duration per session over age. Exposure durations (*t* in minute) are estimated to ensure that the peak fluence (99^th^ percentile of the target^18^) at dlPFC and vmPFC reaches an optimal fluence of 3 J/cm^2^ per session for F3-F4 and Fpz positions, respectively. Note that in our earlier study,^18^ a treatment session is effective when the upper quartile of the target reaches fluence of 3 J/cm^2^, whereas the 99th percentile of the target is used in this paper. The revision is made in order to minimize the light fluence on the skin and reduce the risk of overexposure. Furthermore the engagement by t-PBM of the most superficial cortex, for any given brain area, is considered sufficient to modulate the emotion regulation circuitry. The expression of *t* (in second) can be written as

**Fig 8.**
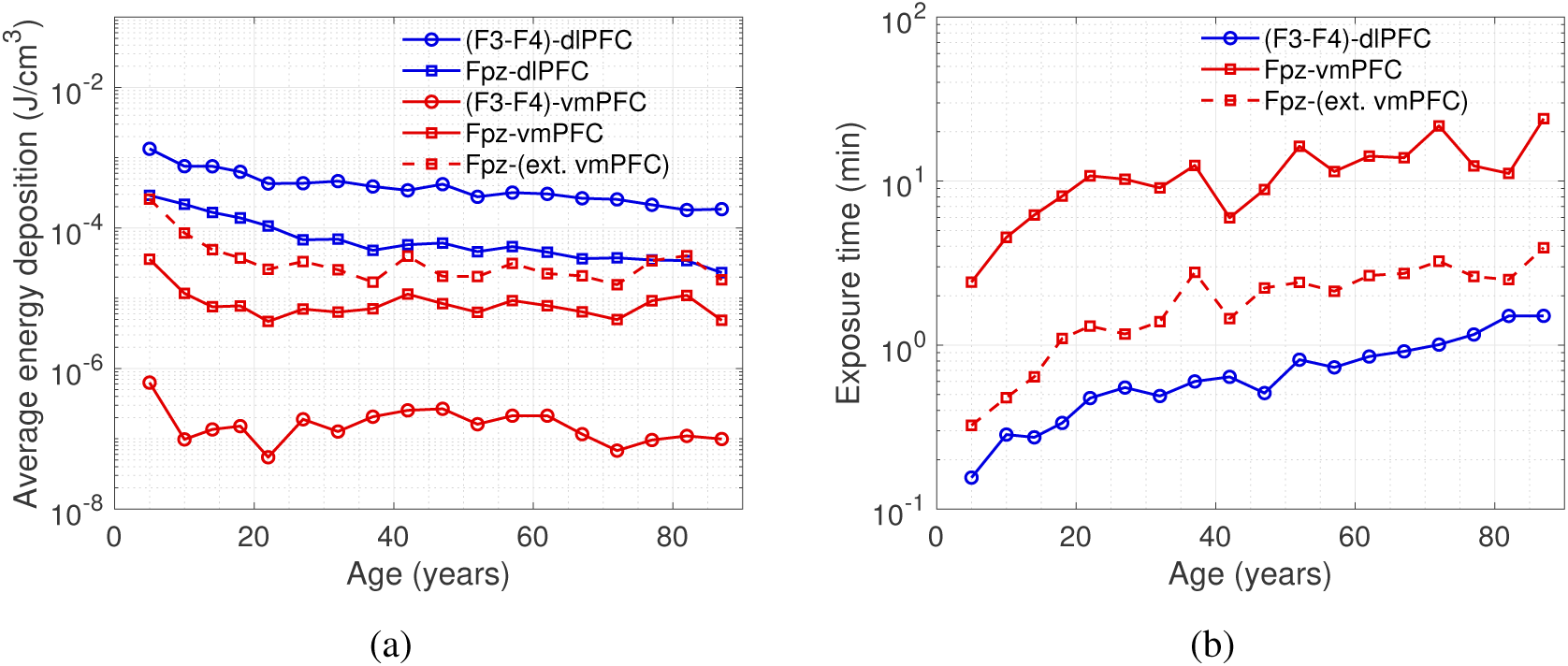
Plots of (a) average energy deposition and (b) estimated treatment duration (in minute) for dlPFC and vmPFC. Each line represents the result for a source-region pair, as listed in the legends. All results are computed at 810 nm. The dashed lines represent the results for extended vmPFC (ext. vmPFC). To estimate exposure time, we assume that the skin irradiance is 300 mW/cm^2^.

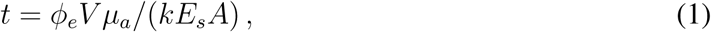

where *ϕ*_*e*_ (J/cm^2^) is the effective fluence achieved at the target region-of-interest (ROI), *V* (cm^3^) is the volume with energy deposition above the 99^th^ percentile inside the ROI, *µ*_*a*_ (cm^*−*1^) is the ROI absorption coefficient, *k* ∈ [0, 1] is the percentage of the source energy delivered to the volume with energy deposition above the 99^th^ percentile inside the ROI, *E*_*s*_ (W/cm^2^) is the skin irradiance of the source, and *A* (cm^2^) is the size of the illumination area. The skin irradiance is set at 300 or 30 mW/cm^2^ for computing the exposure duration. The illumination area is 28.4 cm^2^ and 28.4 × 2 cm^2^ for Fpz and F3-F4 positions, respectively, based on the simulated source device.^18^

In Fig. 9(a), we plot the ECT thicknesses over age groups for F3, F4 and Fpz positions. The correlations between the ECT thickness and the average energy deposition (in log10 scale) are demonstrated in Fig. 9(b) in the target region using 810 nm illumination. The target region is dlPFC for F3/F4 placement and vmPFC/extended vmPFC for Fpz source placement. In our results, due to the low thickness and low absorption, the CSF layer shows minor effects on energy deposition across age compared to the ECT layer. Thus the CSF layer is not considered in Fig. 9.

**Fig 9.**
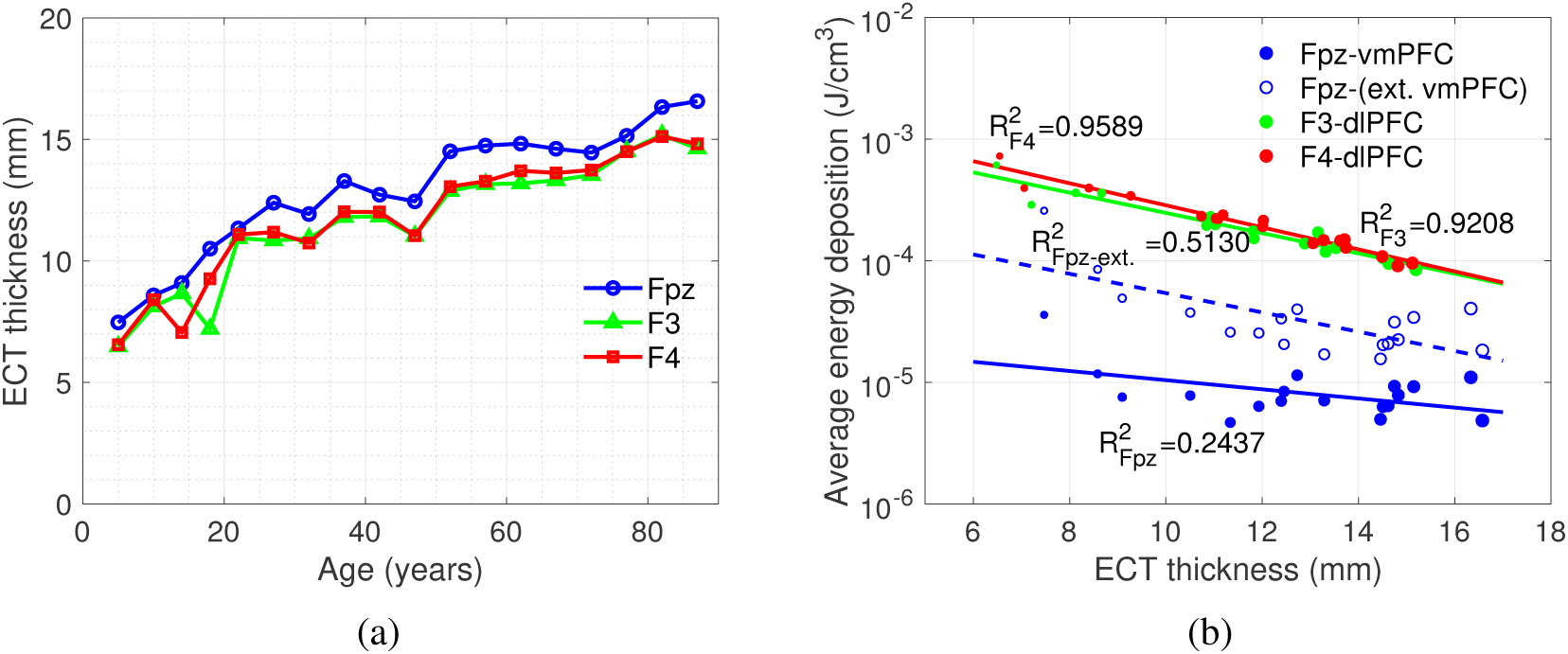
(a) The relationship between age and thickness of extra-cerebral tissues (ECT). (b) The correlation between thickness of ECT and average energy deposition (in log_10_ scale) in the dlPFC and vmPFC. For F3 and F4 positions, the target region is dlPFC, while for Fpz position the target region is vmPFC/extended vmPFC (ext. vmPFC). The size of the markers represents the age group.

## 4 Discussions

From the sample energy deposition maps in Fig. 5, we can visually observe that light penetration to the brain decreases as age increases. The increasing brain size and ECT layer thickness can also be observed in Fig. 5, which impedes energy delivery to the desired brain tissues.

In Fig. 6, all wavelengths show an overall decreasing energy deposition as increasing age for both dlPFC and vmPFC. The linear correlation coefficients between 810 nm and other tested wavelengths are found to be above 0.99 for both ROIs, suggesting that the age-variation has a weak dependency on wavelength. Comparing between wavelengths, 810 nm delivers the highest energy deposition; 850 nm and 1064 nm deliver more energy than 670 nm and 980 nm in most cases. These findings generally agree with previously published simulation-based studies,^18, 58^ however, we want to point out that there is a wide range of brain optical properties in literature, due to diverse measurement techniques and experimental settings. Different choices of literature values could potentially lead to different rankings in wavelength efficiency.^58, 59^

Fig. 7(a) shows that the top-5 regions in energy deposition are mostly located in the frontal lobe of the brain, which also includes the dlPFC. The rostral middle frontal gyrus and the caudal middle frontal gyrus that compose the dlPFC receive approximately 10-fold more energy deposition than the remaining three. This is mainly a result of shorter distances between these two regions and the source when compared to other parcellations. In Fig. 7(b), the energy deposition in the frontal pole region is over 10-fold higher than that of other parcellations. Furthermore, the first 4 parcellations are not part of vmPFC. This is due to the fact that the vmPFC is located at the bottom of cerebral hemispheres, as shown in Fig. 2(b), while the Fpz source position delivers light directly toward the frontal pole. Across all age groups, the F3-F4 position delivers 0.23-2.9% of the total energy into dlPFC while the Fpz position only delivers 0.0066-0.09% of the total energy into vmPFC. Based on this result, the Fpz position appears to be less effective in delivering light to the target region in comparison with the F3-F4 position. In both Figs. 7(a) and 7(b), we can observe a decrease in energy deposition, as age increases, similar to the overall trend shown in Fig. 6.

In Fig. 8(a), decreases in energy deposition can again be seen for all source-region pairs as age increases. However, energy deposition at the vmPFC shows a stronger decrease in younger age groups than that at the dlPFC. For example, the energy deposition decreases by 86.97% and 67.98% from 5 to 20 years old for Fpz-vmPFC and (F3-F4)-dlPFC, respectively. In addition, during adulthood, the vmPFC has a slower decay rate in energy deposition along age than the dlPFC. With frontal pole included, the extended vmPFC has an average 4-fold increase in energy deposition across age groups compared to the vmPFC. Therefore the actual energy deposition at the vmPFC may vary depending on the definition of the target region. Furthermore, the energy deposition of Fpz-dlPFC is in general higher than that of Fpz-vmPFC as well as Fpz-(extended vmPFC) across age, which is caused by the location of vmPFC illustrated previously. This suggests an effective t-PBM treatment targeting at vmPFC/extended vmPFC with an Fpz source also delivers sufficient dosage to dlPFC, but not vice versa.

The plots in Fig. 8(b) show that the desired treatment duration increases with age for both (F3-F4)-dlPFC and Fpz-vmPFC treatments. For (F3-F4)-dlPFC with skin irradiance of 300 mW/cm^2^, the effective fluence can be achieved in less than two minutes across all ages. However, the exposure duration is much longer for Fpz-vmPFC and is around 10 to 20 min after 20 years of age. This is due to insufficient energy deposition at the vmPFC discussed earlier. The addition of frontal pole to the extended vmPFC reduces the exposure duration by 71%-88% compared to the vmPFC since the Fpz position mostly concentrates energy at the frontal pole. Caution should be exercised when applying these findings to clinical research or practice. It is in fact unusual to expose the skin for up to 20 min to high-irradiance of 300 mW/cm^2^. Our data also suggest the potential for overexposure of the most superficial brain areas, such as the frontal poles when the light source is positioned near Fpz. Furthermore, there may be other treatment strategies with different skin irradiance. From Equ. 1, we can see that given a fixed illumination area, the factors that determine the treatment exposure duration are the effective fluence and irradiance. In addition, the exposure duration is inversely proportional to the skin irradiance. Therefore, one can adjust the exposure duration by increasing or decreasing the skin irradiance.

In Fig 9(a), we notice that for all F3, F4 and Fpz positions, the thickness of the ECT layer increases as age increases. The ECT regions under the F3 and F4 positions have very similar thickness values across age due to symmetry. In comparison, the ECT thickness for Fpz position is generally larger than the other two, which coincides with the findings reported previously.^60^

In Fig. 9(b), the plots between the ECT thickness and the log-scaled energy deposition show a rough linear relationship between the two parameters. Applying linear regressions, we obtained four linear models for 1) Fpz-vmPFC: *y* = −0.0376*x* − 4.6061 (*R*^2^=0.2437), 2) Fpz-(extended vmPFC): *y* = −0.0794*x* − 3.4733 (*R*^2^=0.5130), 3) F3-dlPFC: *y* = −0.0833*x* − 2.7749 (*R*^2^=0.9208) and 4) F4-dlPFC: *y* = −0.0906*x* − 2.6388 (*R*^2^=0.9589). For both F3 and F4 positions, the ECT thicknesses show a strong negative linear correlation to the average energy deposition in log-scale at the dlPFC. The plots in Fig. 9(b) show discernible deviations from a linear-fit in younger ages, possible due to the boundary effect in smaller-sized head models. Only a weak linear correlation is found for Fpz position. It is our belief that the relatively weak correlation at the Fpz source is a result of larger separation between the target region (vmPFC) and the source. Nonetheless, an overall decreasing trend is evident for the vmPFC energy deposition from the Fpz-centered source. For Fpz sources targeting the extended vmPFC, the result presents a stronger linear correlation compared to Fpz-vmPFC. The frontal pole merged within the extended vmPFC is closer to the Fpz position and shows a strong linear correlation (*R*^2^=0.8660), which raises the overall correlation for the Fpz-(extended vmPFC). These results indicate that the anatomical development of the head, and especially the increase of ECT thickness, is largely responsible for decreased energy deposition in the brain as a result of growth and aging. Furthermore, shorter distances between the source position and target region results in greater correlation between energy deposition and ECT thickness.

In addition, we have repeated our simulations using refined brain anatomical models by further separating the ECT layer into the scalp and skull layers using FSL. Our simulations using such atlases show very similar results (not included) to our above findings. This suggests that the overall ECT tissue thickness plays a more important role in t-PBM treatment compared to the individual scalp/skull thicknesses.

## 5 Conclusion and summary

In this report, we systematically investigated light dosimetry in t-PBM treatment across a wide span of ages. To ensure accurate quantification, we used anatomically appropriate brain atlases and state-of-the-art Monte Carlo simulations. For modeling the brain anatomy, we have developed a robust workflow to create multi-layered head segmentations from publicly available neurodevelopmental MRI atlas library data. For each brain segmentation, we have also generated corresponding brain parcellations to facilitate quantitative analysis. A number of conclusions have been found from our simulation results, and many of those are consistent with our previous study.^18^ First, the energy deposition decreases across lifespan regardless of the source position and parcellations. However, the decline is faster before adulthood and this is more noticeable for the vmPFC. Secondly, wavelength selection shows a negligible impact to the trend of energy deposition decay over age, despite that the 810 nm consistently gives the highest energy deposition compared to 4 other commonly used wavelengths. Although this result is generally consistent with previous simulation-based studies, ^13, 61^ we would like to highlight that our results are dependent on the choices of brain optical properties. As we discussed in,^18^ there is no widely accepted set of optical properties for brain tissues. Using values from different literature may lead to different preferences in wavelength.^59^ Thirdly, a negative linear correlation is observed between the thickness of ECT layer and the log-scale average energy deposition in two selected brain ROIs, suggesting that the brain anatomical changes are largely responsible for the observed age-dependent variations.

Furthermore, the Fpz source position has longer separation distances from the vmPFC region compared to the separations between the dlPFC and the F3-F4 source, suggesting that higher source intensity (alternatively, longer exposure time) is required when targeting the vmPFC region compared to the dlPFC region. Also, the frontal pole (#6 in Fig. 2(d)) shows stronger energy deposition compared to the remaining parts of vmPFC. Thus the exposure duration for the extended vmPFC (vmPFC+frontal pole) is shortened. In addition, we provide a simple approach to estimate the treatment duration using our simulated results. From this relationship, the exposure duration in general increases as age increases. These quantitative assessments are expected to provide clinicians guidance in designing personalized PBM treatment plans and ultimately improve the outcome of the procedures.

We would like to mention that there are several known limitations in this study. First, the simulations are performed using averaged brain atlases derived from public datasets. We acknowledge that using subject-specific brain segmentations may lead to more accurate estimations. In fact, our reported processing methods can be directly applied to subject-specific anatomical scans if such data become available, making it suitable for personalized t-PBM treatment planning. Secondly, the brain optical properties are assumed to be static across age groups. While age-dependent brain tissue optical property measurements are generally lacking, earlier studies on breast tissues have observed little or no correlation with age.^62, 63^ A correlation between dermis tissue absorption over age was reported,^64, 65^ but the magnitude of such variation is relatively small. Our analyses can be easily extended to include age-dependent optical properties when such data become available in the future. Thirdly, this study is specifically focused on treating MDD by performing t-PBM to target vmPFC and dlPFC brain regions. If modeling other t-PBM source forms and brain ROIs becomes necessary, we can apply the same multi-layered brain segmentation and parcellation models to quickly recreate the results for the new targeted regions. Lastly, we largely rely on dedicated brain segmentation tools, namely, FSL and FreeSurfer, to create the brain anatomical models used in this study. The choice of neuroanatomical analysis tools may lead to variations in brain segmentations and simulation results. In the next steps, we may extend this study to use more accurate mesh-based Monte Carlo^66, 67^ simulations and subject-specific scans.

## Acknowledgments

This work is supported by the National Institute of Health (NIH) under grants R01-GM114365, R01-CA204443 and R01-EB026998. We would like to thank Dr. Lilla Zöllei for her inputs regarding FreeSurfer software, and Edward Xu for his help on manuscript development.

**Yaoshen Yuan** is a doctoral candidate at Northeastern University. He received his BE degree from Southeast University, China, in 2014 and MSE from Tufts University in 2016. His research interests include Monte Carlo simulation for photon transport, GPU algorithm enhancement and signal processing.

**Paolo Cassano**, MD, PhD, is a principal investigator at the Center for Anxiety and Traumatic Stress Disorders and at the Depression Clinical and Research Program, Massachusetts General Hospital; and an assistant professor at Harvard Medical School. He specializes in photomodulation for psychiatric disorders. In 2015, his work on photomodulation was featured by the Boston Globe as one of the breakthroughs for the toughest medical conditions. He was also interviewed by CNN and by the Washington Post.

**Matthew Pias** is a graduate student at Northeastern University. He received his MS degree from Northeastern University in 2019 and his BSE from the University of Connecticut in 2017. His research interests are in image processing and biomedical optics.

**Qianqian Fang**, PhD, is currently an assistant professor in the Bioengineering Department, North-eastern University, Boston, USA. He received his PhD degree from Dartmouth College in 2005. He then joined Massachusetts General Hospital and became an assistant professor in 2012, before he joined Northeastern University in 2015. His research interests include translational medical imaging systems, low-cost point-of-care devices for resource-limited regions, and high performance computing tools to facilitate the development of next-generation imaging platforms.

